# *μ*-Opioid Endomorphins and DDP-IV Inhibitor Sitagliptin Enhance Amyloid-Beta Clearance and Memory in an Alzheimer’s Cell Model

**DOI:** 10.1101/2025.07.24.666703

**Authors:** Maxx Yung, Wei Zhu

## Abstract

Alzheimer’s Disease is a neurodegenerative disorder caused by A*β*_42_ aggregation. Endomorphins 1 and 2 (EM1, EM2), two novel *μ*-opioid agonists, have been implicated in protecting against A*β*_42_ toxicity, though it is unclear how the endomorphins achieve their effects. Phase one of the study found that EM1 and EM2 activation protected A*β*_42_-treated cells. This protection, mediated by *μ*-opioid receptor (MOR) activation, also reduced rotenone-induced oxidative stress, both in a dose-dependent manner.

Pretreatment with naloxone, a *μ*-opioid antagonist, reversed these effects, confirming MOR involvement in EM1 and EM2’s actions. In phase two, molecular docking techniques suggested that sitagliptin can prevent intracellular EM1 degradation. In vitro assays demonstrated that sitagliptin enhanced intracellular EM1’s beneficial effects in promoting cell survival and reducing cell apoptotic activity, A*β*_42_ aggregation, and hydrogen peroxide free radical concentrations. This suggests intracellular EM1 can mitigate the toxic effects of A*β*_42_ aggregation. However, sitagliptin did not enhance EM1’s effects on BDNF expression or neurite outgrowth, suggesting that MOR activation, rather than intracellular EM1, primarily drives mechanisms associated with memory improvement. Collectively, our findings suggest that both intracellular EM1 and EM1-mediated MOR activation offer potential therapeutic avenues for mitigating memory impairment in Alzheimer’s and potentially COVID-19. Furthermore, this research underscores the critical role of the MOR in broader memory mechanisms.

## Introduction

### Alzheimer’s Disease

Alzheimer’s disease (AD), a neurodegenerative disorder characterized by progressive memory impairment, accounts for 60-75% of the 55 million dementia cases globally [1, 2]. In the United States, AD ranks as the sixth leading cause of death [3]. The risk of developing AD increases dramatically with age—from approximately 3% at age 65 to nearly 50% by age 85 [2]. With projected rises in life expectancy, both AD prevalence and associated care costs are expected to increase substantially [4], potentially doubling the economic burden of patient care [5]. In the United States alone, prevalence is projected to rise from 5 million in 2007 to 13 million by 2050 [2]. These alarming trends emphasize the urgent need for effective AD treatments and a better understanding of its impact on memory processes. This study addresses this need by investigating the potential of endogenous opioids to mitigate the damaging effects of amyloid beta (A*β*) plaques, a key pathological hallmark of AD.

The agglomeration of A*β* monomers into A*β* plaques is believed to be the principal cause of AD [2, 6–11]. In the AD brain, the overproduction of abnormally long A*β*_42_ monomers leads to increased aggregation of A*β*_42_ plaques, particularly in the hippocampus [12]. This region of the brain critical for spatial learning, memory processing, long term potentiation (LTP), and synaptic plasticity [13–18]. This results in neuronal cytotoxicity, death, and thus memory impairment [19, 20]. Additionally, A*β*_42_ aggregation is believed to contribute to extensive oxidative damage [21–29]. Thus, reducing A*β*_42_ induced cellular damage and death is the primary focus of current therapeutic approaches to AD.

### The *μ*–Opioid Receptor

Emerging research suggests using opioids as a potential treatment for AD [7]. Opioid receptors and their corresponding peptides are expressed endogenously in the central nervous system [18, 30–32] and are widely distributed in hippocampal CA1, CA2, and CA3 regions [7, 14, 33–35]. Of particular interest is the *μ*-opioid receptor (MOR) [32, 33, 36], as previous work on MOR has demonstrated that MOR concentrations in the hippocampus decreased with increasing age [37–40]. Interestingly, a prior study found that the hippocampus, exhibiting the highest aggregation of A*β*_42_, suffers the most damage in elderly AD patients [2].

Additional studies have shown that the MOR modulates memory processes, cognition, and learning, along with its well-documented impacts on nociception [7, 37, 41–49].

While the MOR’s role in memory, learning, and cognition is established, the direction of its influence – whether activation is beneficial or detrimental – remains debated. In the context of AD, some studies suggest potential benefits: for instance, Shiigi, Takahashi, and Kaneto [50] and Cai and Ratka [7] found that MOR activation may facilitate memory retrieval in mice. Conversely, Jang et al. [51] reported that MOR knockout mice exhibited spatial memory deficits and reduced hippocampal LTP, suggesting a requirement for MOR signaling in normal memory function. This complex picture highlights the need for further investigation into the role of the MOR in AD and its potential as a therapeutic target.

### Rationale

This study will assess how MOR activation affects the outcome of A*β*_42_ induced cell death in SK-N-SH neuronal cells. MOR agonists endomorphin-1 (EM1, Tyr-Pro-Trp-Phe-NH2) and endomorphin-2 (EM2, Tyr-Pro-Phe-Phe-NH2) were used to selectively activate the MOR. These mu-opioid agonists demonstrate extremely high selectivity for the MOR receptor over the DOR and KOR opioid receptors [52].

Interestingly, the distribution of endogenous opioids in the human brain is inversely correlated with retgions of A*β* aggregation [3, 53]. This inverse relationship suggests potential inhibitory interactions. Therefore, we hypothesized that prolonging the half-lives of EM1 and EM2, by inhibiting the dipeptidyl peptidase IV (DPP4), could influence A*β* aggregation. Previous studies showed that DPP4 degrades various brain neuropeptides [5, 43, 54–60]. Inhibiting DPP4 may consequently prevent the degradation of *μ*-opioid agonists, extending the biological activity of intracellular EM1 and EM2.

Furthermore, we sought to investigate the impact of EM1 and EM2 on hippocampal memory-related processes. To this end, we examined neurite outgrowth and brain-derived neurotrophic factor (BDNF) levels. Neurite outgrowth is crucial for neuronal connectivity, with its disruption being a hallmark of neurodegeneration [61, 62]. BDNF, in turn, plays a vital role in memory and learning, including promoting neurite outgrowth itself [49, 63–65], making these key indicators of neuronal health and memory function.

## Materials and Methods

### Chemicals

MOR-specific antagonist naloxone (NX), MOR-specific agonists EM1 and EM2, and the DPP4 inhibitor sitagliptin were purchased from Cayman Chemicals. EM1 and EM2 were serially diluted to achieve final concentrations of 20 μM, 5 μM, 1 μM, and 0.1 μM. NX and sitagliptin were diluted to a 1 μM concentration.

Human A*β*_42_ peptides (pre-aggregated oligomeric form) and rotenone were purchased from Sigma Aldrich. A*β*_42_ oligomers were reconstituted to a final concentration of 10 μM in sterile PBS (pH 7.4) buffer, and rotenone was prepared at 1 μM concentration in DMSO. The oligomeric state of A*β*_42_ was used without further manipulation as confirmed by the manufacturer’s specifications.

### Cell Culturing

SK-N-SH neuronal cells were obtained from the American Type Culture Collection (ATCC). The SK-N-SH cells were cultured with Eagle’s Minimum Essential Medium and supplemented with penicillin and fetal bovine serum at 5% concentration. Cells were grown in an incubator at 5% CO_2_ and 37°C. Cells were then collected and distributed into 6-well plates or 96-well plates at an average seeding density of 200000 cells per well and 5000 cells per well, respectively.

### Molecular Docking

Molecular docking was utilized to predict whether EM1 would be cleaved by DPP4 [66–68]. Both EM1 and sitagliptin (DPP4 inhibitor) were docked to the active site of a human recombinant DPP4 structure. We then compared the docking position, orientation, and conformation of EM1 to that of sitagliptin. 3D protein and ligand structures of EM1, sitagliptin, and DPP4 were obtained from the PubChem and RCSB Protein Bank databases. Prior to docking, all water molecules were removed from the protein structure, and ligands underwent energy minimization to ensure optimal conformations. PyRx (version 0.8) running AutoDock Vina [69] was used to simulate the protein-ligand interactions following the procedures of Dallakyan [70]. For each ligand, 10 independent docking runs were performed to ensure thorough sampling of possible binding modes. Binding energies below -9.0 kcal/mol generally indicate strong interactions, while those approaching -10.0 kcal/mol suggest very high binding affinity. Protein-to-ligand interaction models were generated and visualized using BioVia Discovery Studio Visualizer (2021 version).

### Assays

Cell survival rates were assessed using MTT assays, following the method of van Meerloo et al. [71]. Cells were seeded in a 96-well plate and incubated for 24 hours post-treatment. Optical density was measured at 595 nm using a microplate absorbance reader. Samples were assayed in triplicate. The results were expressed as a percentage of the control, which was set at 100%.

Caspase–3/7 and hydrogen peroxide (*H*_2_*O*_2_) assays were performed to quantify apoptosis activity [72] and free radical concentrations, which contribute to oxidative damage [73]. 10 μL treatments were added into 6-well plates containing cells and incubated for 24 hours. For the caspase–3/7 assay, cells were then collected and transferred into 96-well plates and 50 μL of caspase assay buffer, 45 μL of LDH, and 5 μL of caspase substrate were added into each well. Optical density was measured at 415 nm in a microplate absorbance reader at 15-minute intervals. The colorimetric *H*_2_*O*_2_ assay kit was purchased from Assaygenie, and the provided instructions were followed.

Neurite outgrowth assays were performed to measure neuronal connectivity.

Following the addition of 10 μL of treatments to 6-well plates for 24 hours, the media were aspirated, and the cells were stained using a standard HEMA-3 staining protocol [74]. Neurons were imaged under a light microscope, and neurite intersections and lengths were analyzed using the NeuronJ and Simple Neurite Tracer plugins on the freely available Fiji ImageJ software (version 1.53).

Human BDNF and A*β*_42_ double-antibody sandwich ELISA kits were purchased from Invitrogen and the provided instructions for a standard double-antibody sandwich ELISA protocol were followed. Optical density was measured at 450 nm in a microplate absorbance reader.

### Statistical Analysis

GraphPad Prism 9 software (version 9.4.1) was used for all statistical analyses.

Two-way multiple ANOVAs followed by Bonferroni *post hoc* tests were used to compare the differences between all pairs of conditions. The significance for statistical analysis was set at *p* < .05.

## 1 Results

### 1.1 EM1 and EM2 Protect Against A*β*_42_ Induced Cell Death

In healthy SK-N-SH cells, increasing concentrations of EM1 and EM2 treatment decreased cell survival rates compared to control (*p* < .001, Fig. 1a). Conversely, in A*β*_42_ treated cells, increasing concentrations of EM1 and EM2 treatment increased cell survival rates compared to A*β*_42_ treated cells without EM1 or EM2 treatment (*p* < .001, Fig. 1b).

**Figure 1a:**
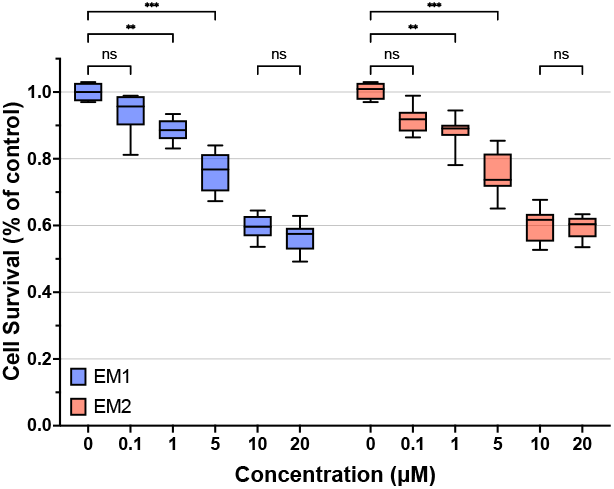
EM1 and EM2 Treatment in Healthy SK-N-SH Neuronal Cells (*F*_5,91_ = 221.5)

**Figure 1b:**
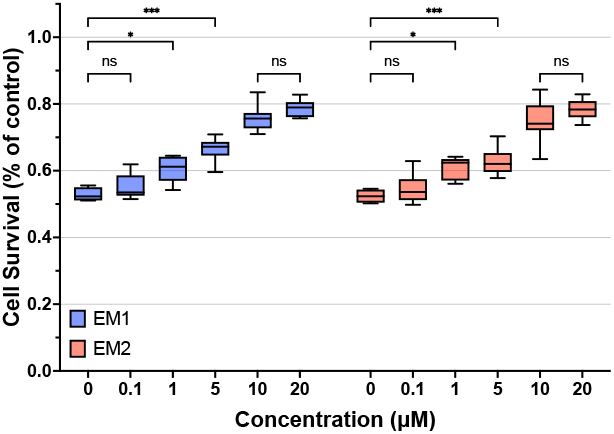
EM1 and EM2 Treatment in A*β*_42_ Treated SK-N-SH Neuronal Cells (*F*_5,94_ = 118.8)

In healthy cells, inhibition of the MOR through NX treatment significantly lowered cell survival. In cells exposed to A*β*_42_, NX pretreatment negated any increase in cell survival due to EM1 and led to lowered survival rates compared to EM1 treatment (*p <* .001, Fig. 2).

**Figure 2:**
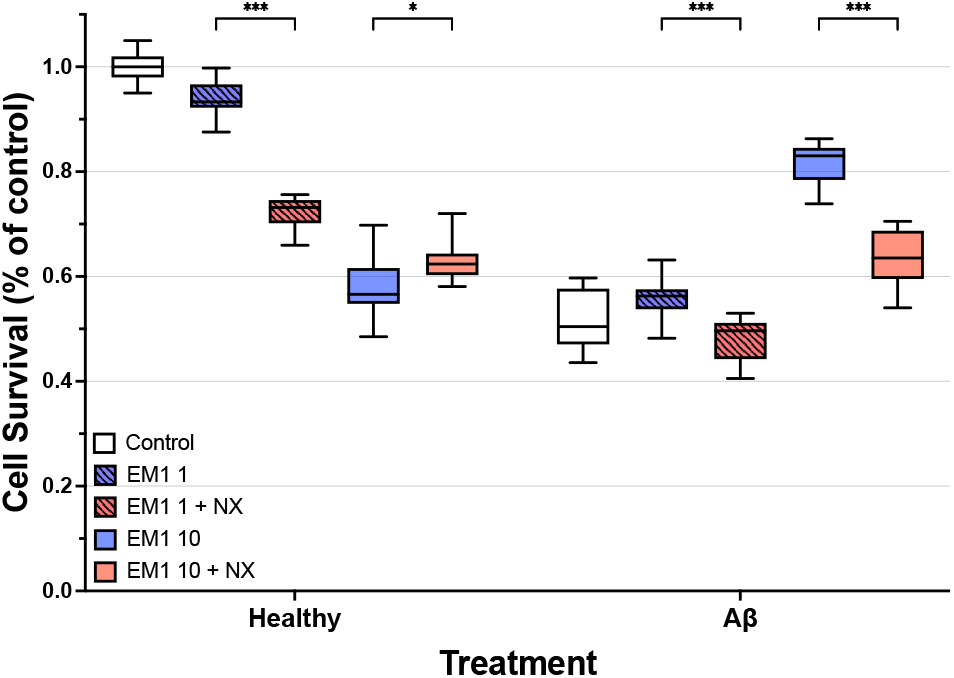
Effects of NX Pretreatment in A*β*_42_ Treated SK-N-SH Cells (*F*_5,172_ = 286.3)

### 1.2 EM1 and EM2 Protect Against LPS Induced Neuroinflammation and Rotenone Induced Oxidative Stress

The addition of 1 μM or 10 μM of EM1 or EM2 had no significant effect on LPS-treated cells (*p* > .05, Fig. 3). Similarly, the addition of 1 μM of EM1 or EM2 had no significant effect on rotenone treated cells. However, the addition of 10 μM of EM1 or EM2 significantly increased survival rates in rotenone-treated cells (*p* < .001, Fig. 4).

**Figure 3:**
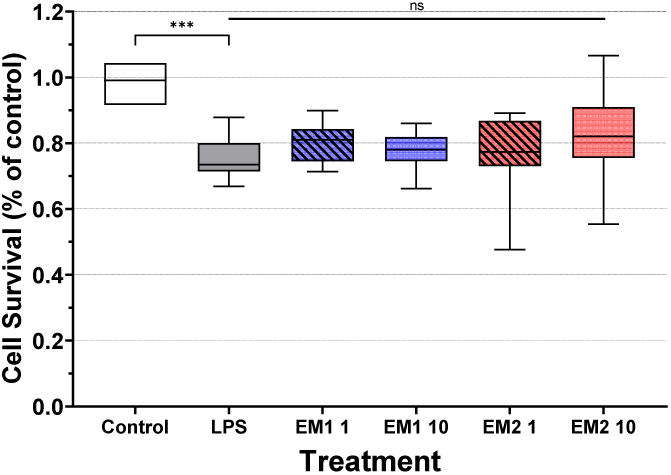
EM1 and EM2 Treatment on LPS-induced Neuroinflammation (*F*_5,88_ = 19.00)

**Figure 4:**
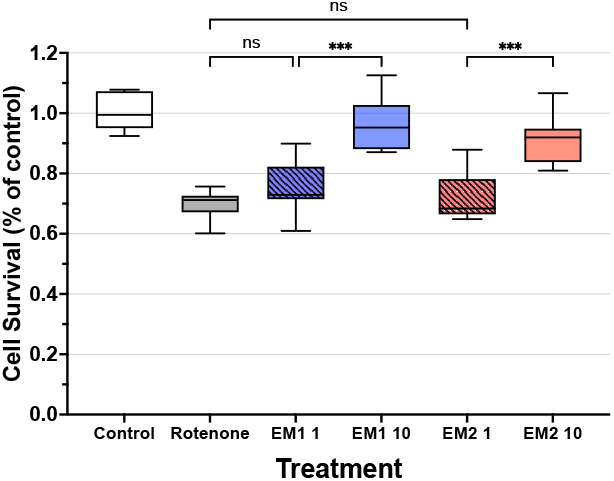
EM1 and EM2 Treatment on Rotenone-induced Oxidative Stress (*F*_5,73_ = 59.03)

## 2 Results

### 2.1 High Binding Affinity of EM1 With the DPP4 Binding Domain

Molecular docking of EM1 and NX with the DPP4 binding domain yielded high binding energies of -9.4 kcal/mol and -9.1 kcal/mol at high accuracies (RMSD ≤ 1.5 Å) (Table1). For comparison, docking EM1 and NX to their endogenous MOR binding domain resulted in even higher binding energies of -10.7 kcal/mol and -10.3 kcal/mol. The DPP4 antagonist sitagliptin exhibited the highest binding affinity for DPP4, with a binding energy of -11.5 kcal/mol and RMSD 1≤ Å (Table 1).

**Table 1:** Summarized Results of Molecular Docking

| Ligand | Binding Affinity | RMSD | Type of Interaction | Residue Information |
| --- | --- | --- | --- | --- |
| Endomorphin-1 | -9.4kcal/mol | <1.5Å | Hydrogen Bond<br>Pi-Pi Interaction | Phe357, Glu205, Ser209, Tyr662, Tyr585<br>Tyr666, Ser630, Arg125, Tyr547 |
| Naloxone | -9.1kcal/mol | <1.5Å | Hydrogen Bond<br>Pi-Alkyl Interaction | Glu205, Glu206, Tyr662, Tyr666 |
| Sitagliptin | -11.5kcal/mol | <1Å | Hydrogen Bond<br>Pi-Alkyl Interaction<br>Electrostatic Interaction | Tyr128, Val155, Leu115, Ser106<br>Thr156, Trp62, Arg61, Tyr105, Ile107<br>Tyr132, Glu117, Asp104 |

Figures 5a and 5b illustrate that both EM1 and sitagliptin bind to the same DPP4 binding domain (Figs. 5a, 5b). Predicted two-dimensional protein-to-ligand interactions for EM1 and sitagliptin are also shown (Figs. 6a, 6b), detailing the specific types and number of bonds formed between DPP4 and EM1 or sitagliptin. Docking results are summarized in Table 1.

**Figure 5a:**
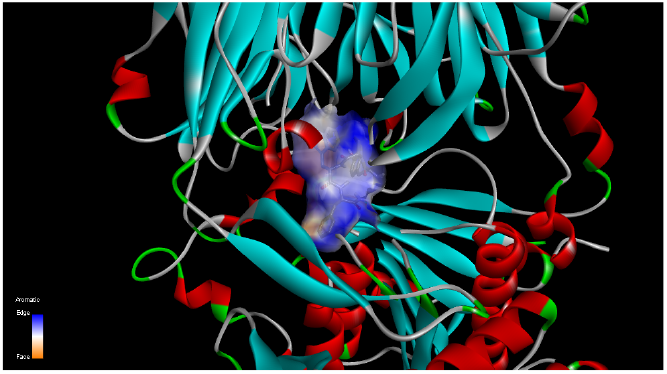
Molecular Docking of EM1 and DPP4

**Figure 5b:**
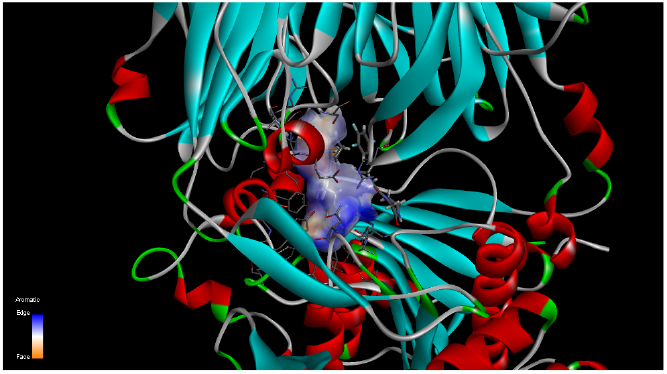
Molecular Docking of Sitagliptin and DPP4

**Figure 6a:**
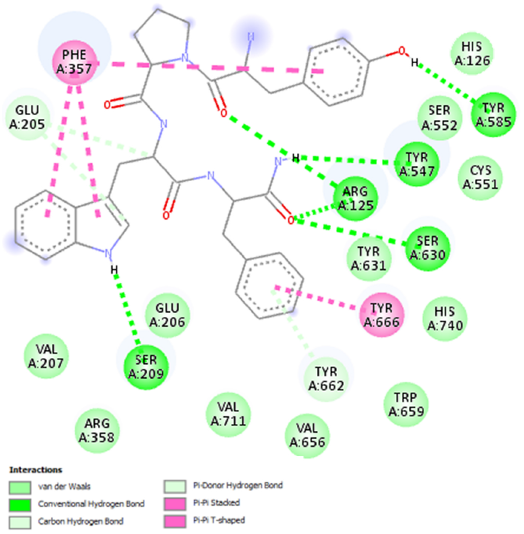
DPP4-EM1 2D Protein-Ligand Interactions

**Figure 6b:**
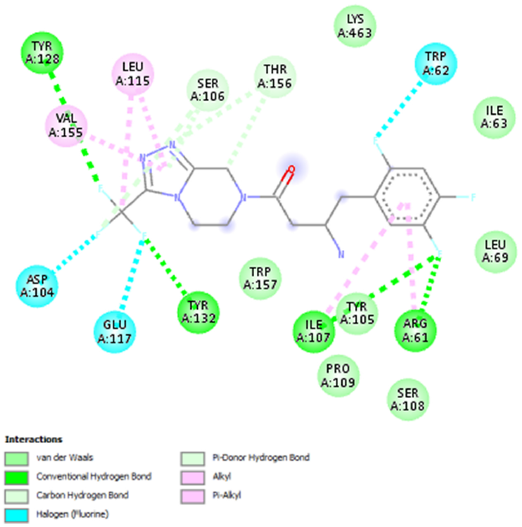
DPP4-Sitagliptin 2D Protein-Ligand Interactions

### 2.2 DPP4 Inhibitor Sitagliptin Enhances Protective Effects of EM1 and EM2

Sitagliptin-mediated DPP4 inhibition combined with endomorphin treatments reduced cell survival rates in healthy SK-N-SH cells compared to endomorphin treatment alone (*p* < .05, Fig. 7a).

**Figure 7a:**
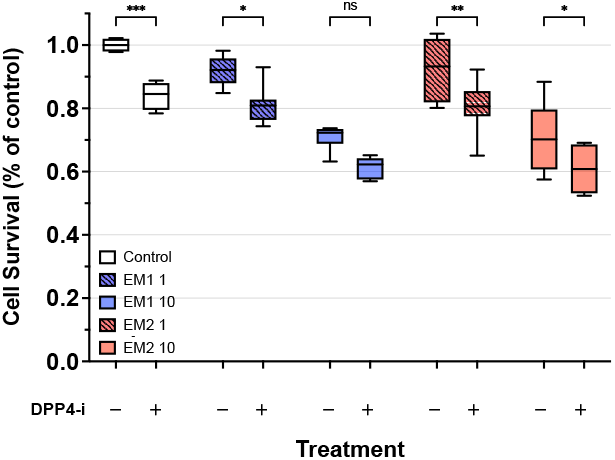
Effects of DPP4 Inhibition on EM1 and EM2 in Healthy SK-N-SH Cells (*F*_4,76_ = 63.78)

**Figure 7b:**
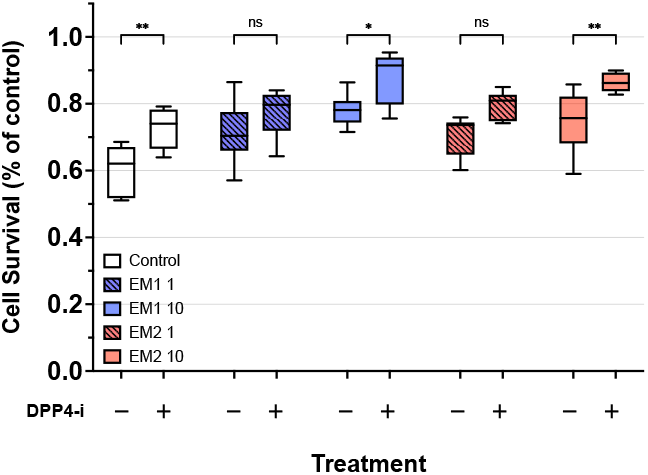
Effects of DPP4 Inhibition on EM1 and EM2 in SK-N-SH Cells Treated with A*β*_42_ (*F*_4,84_ = 17.60)

In stark contrast, when cells were challenged with A*β*_42_, this same combination of sitagliptin and endomorphins significantly increased cell survival rates compared to endomorphin treatment alone (*p* < .001, Fig. 7b). This divergent effect between healthy and A*β*_42_-exposed cells further supports the context-dependent nature of endomorphin activity observed in our earlier experiments.

Overall, 10 μM of EM1 treatment lowered A*β*_42_ peptide concentrations across all treatment groups (*p* < .001, Fig. 8b). Sitagliptin and EM1 treatment resulted in 92.32pg/mL of A*β*_42_ while EM1 treatment alone increased A*β*_42_ concentrations to 122.95pg/mL. Interestingly, NX pretreatment had no significant effect on A*β*_42_ concentration compared to EM1 with and without sitagliptin treatment.

**Figure 8a:**
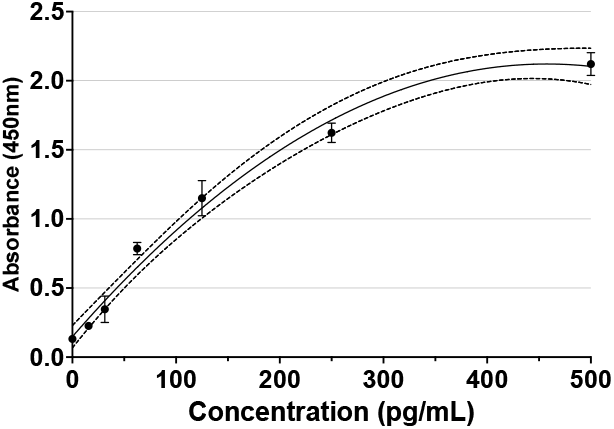
A*β*_42_ Standard Curve

**Figure 8b:**
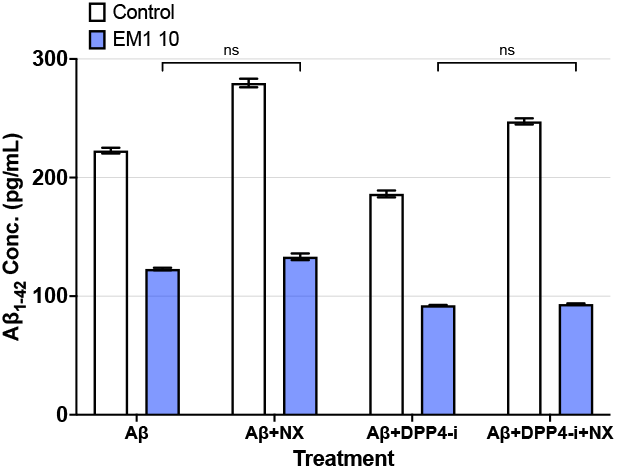
Effect of Sitagliptin and EM1 on A*β*_42_ (*F*_3,8_ = 178.8)

5 μM of EM1 treatment with sitagliptin reduced *H*_2_*O*_2_ concentrations in cells treated with A*β*_42_ by 26.59% compared to EM1 treatment alone (*p* = 0.17, Fig. 9); NX pretreatment reversed this effect increased *H*_2_*O*_2_ concentrations by 52.43%. No significant differences in *H*_2_*O*_2_ concentrations were observed between EM1 and EM2 treatments in the different groups.

**Figure 9:**
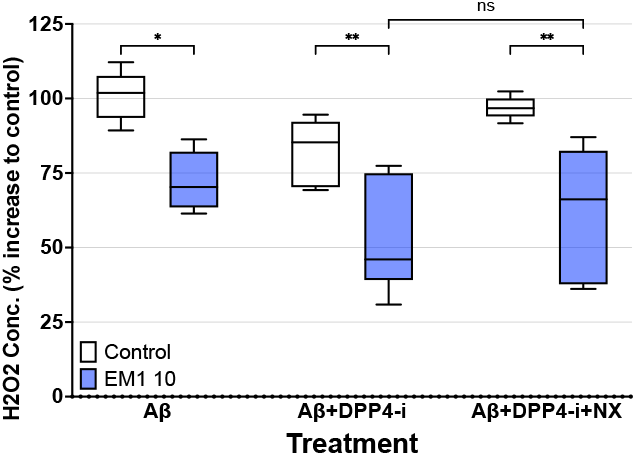
Effect of DPP4 Inhibition and EM1 Treatment on *H*_2_*O*_2_ Free Radical Concentrations (*F*_2,30_ = 6.030)

**Figure 10:**
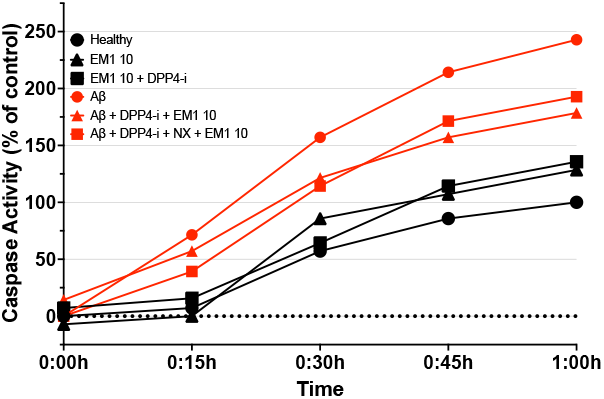
Effect of DPP4 Inhibition and EM1 Treatment on Caspase Activity

Apoptosis rate was indirectly measured by monitoring caspase–3/7 activity. Caspase activity of the control group was normalized to 100%. 5 μM of EM1 with and without sitagliptin treatment slightly increased caspase activity in healthy SK-N-SH cells to 128.57% and 135.71%. Comparatively, A*β*_42_ treatment significantly increased caspase activity to 242.86% compared to control, but adding 5 μM of EM1 with sitagliptin reduced caspase activity by 64.29% compared to A*β*_42_ treated cells. Surprisingly, NX pretreatment also reduced caspase activity by 42.86% compared to A*β*_42_ treated cells.

### 2.3 EM1 and Sitagliptin Treatment Promote Neurite Outgrowth, Improves Neuron Morphology, and BDNF Concentration

Neurons were exposed to different treatment conditions and imaged at 50x magnification. In Fiji ImageJ, images were converted to 8-bit (Fig. 11a) before NeuronJ was used to generate neurite tracings based on color density and edge detection (Fig. 11b). Neurite traces were then isolated and color-coded based on the distance from the soma (Fig. 11c). The SNT neuronal quantification framework in ImageJ was used to perform a linear sholl analysis, which measured neurite trace intersections (Fig. 11d). Greater neurite intersections closer to the soma demonstrate low neurite outgrowth and thus inhibit LTP processes and vice versa.

**Figure 11a:**
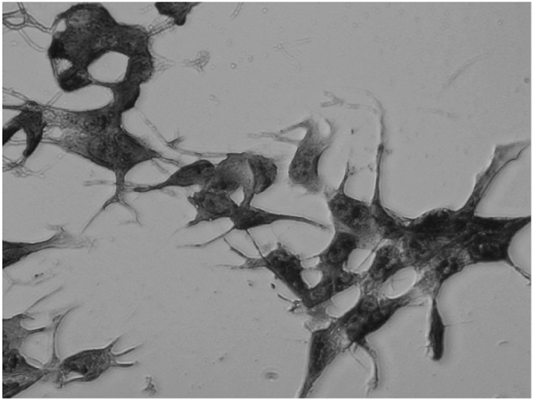
Image of SK-N-SH Neurons under a Microscope

**Figure 11b:**
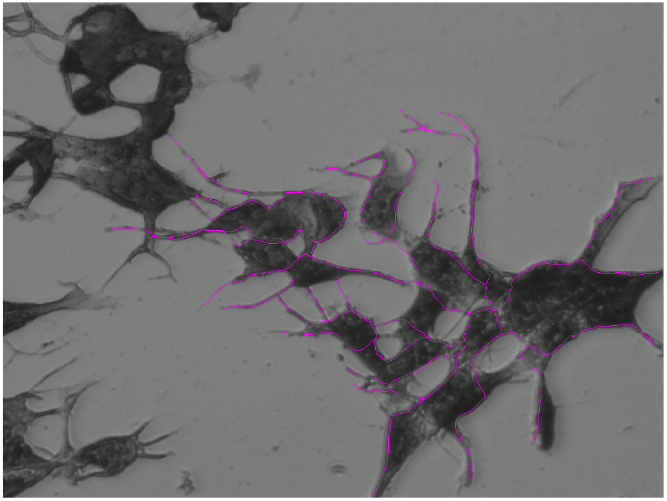
Neurite Tracing using NeuronJ

**Figure 11c:**
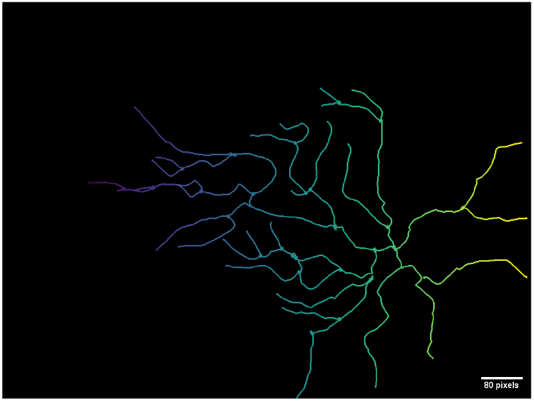
Color-coded Layer Isolation of Neurite Traces using ImageJ

**Figure 11d:**
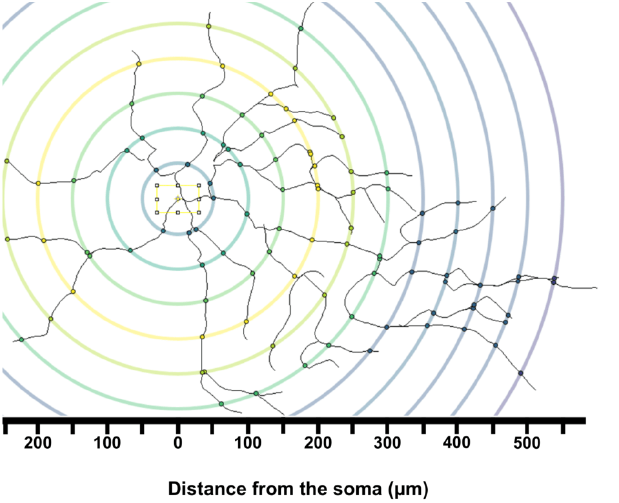
Sholl Analysis of a Sample Neurite Trace using SNT

The addition of 10 μM of EM1 treatment decreased the total number of neurite intersections, especially the number of intersections that occurred further from the soma (Fig. 12a). The addition of 10 μM of EM1 treatment increased the overall number of overall neurite intersections compared to cells only exposed to A*β*_42_ (Fig. 12b). The addition of 10 μM of EM1 and sitagliptin treatment also increased the overall number of neurite intersections compared to cells only exposed to A*β*_42_ and sitagliptin (Fig. 12c). Finally, NX pretreatment did not affect the total number of overall neurite intersections (Fig. 12d).

**Figure 12a:**
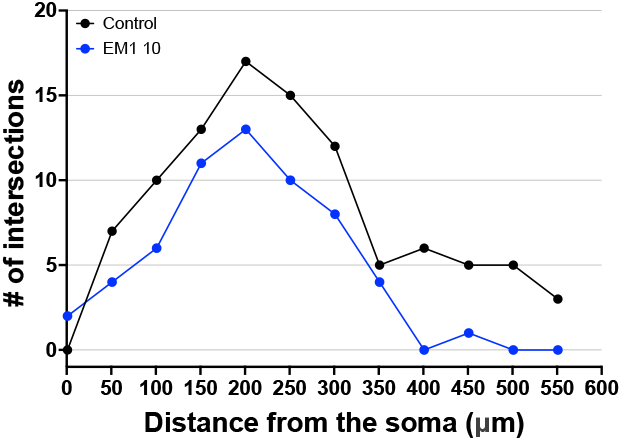
Sholl Analysis of Control Compared to A*β*_42_

**Figure 12b:**
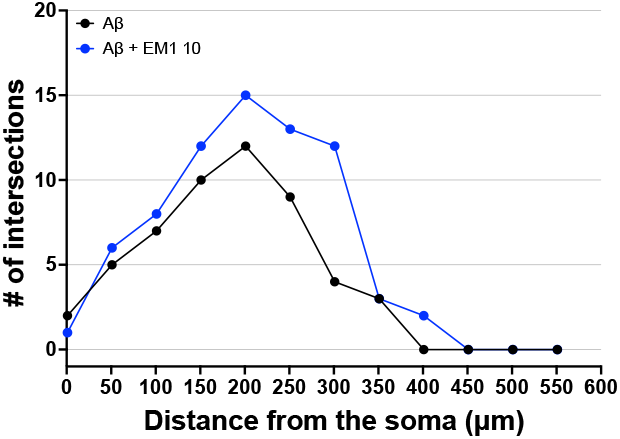
Sholl Analysis of EM1, Sitagliptin, and A*β*_42_ Treatments

**Figure 12c:**
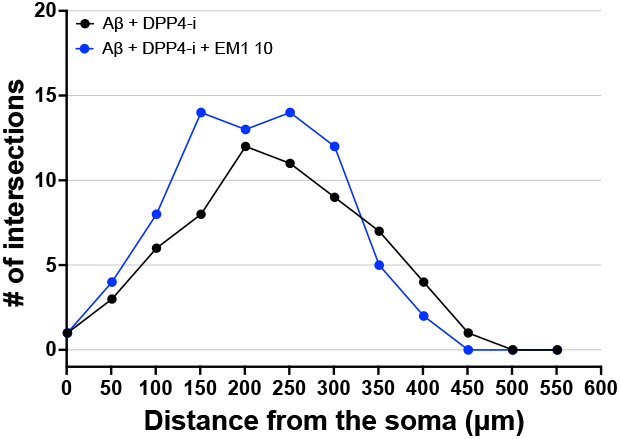
Sholl Analysis of Control Compared to A*β*_42_

**Figure 12d:**
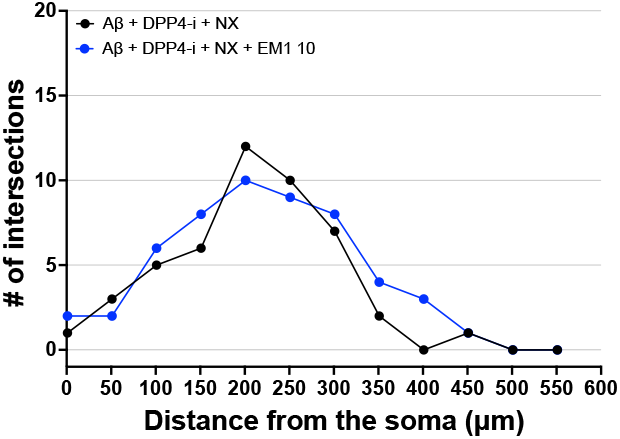
Sholl Analysis of EM1, Sitagliptin, and A*β*_42_ Treatments

Total neurite lengths were also measured across all treatments (*p* < .001, Fig. 13). 10 μM of EM1 and sitagliptin treatment increased total neurite outgrowth compared to only 10 μM of EM1 treatment in cells exposed to A*β*_42_ (*p* < .05). NX pretreatment had no significant effect on total neurite outgrowth.

**Figure 13:**
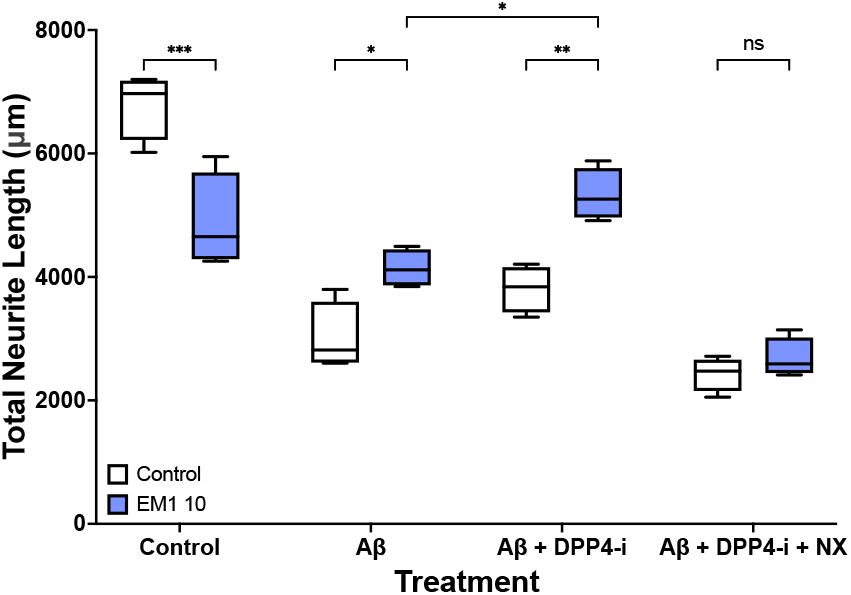
Sholl Analysis of a Neurite Trace (*F*_3,24_ = 21.11)

Consistent with previous results, 10 μM of EM1 treatment decreased BDNF protein concentrations to 664pg/mL from 889pg/mL in healthy cells but increased BDNF protein concentrations to 468pg/mL from 284pg/mL in cells exposed to A*β*_42_ (*p <* .001, Fig. 14b). Additionally, EM1 with sitagliptin treatment further increased BDNF protein concentrations to 677pg/mL from 387pg/mL compared to cells treated with A*β*_42_ (*p* < .001). NX pretreatment had no significant effect on BDNF protein concentration.

**Figure 14a:**
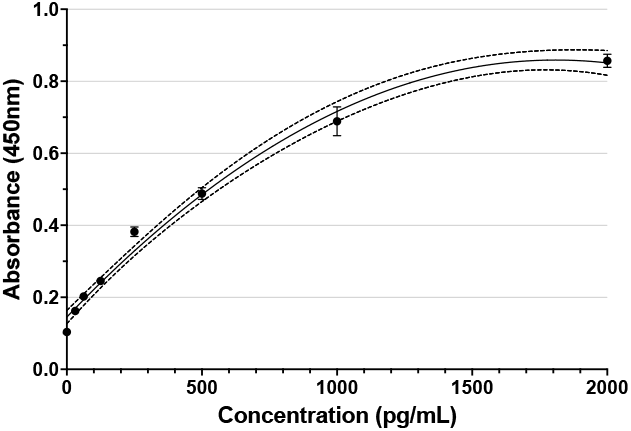
BDNF Standard Curve

**Figure 14b:**
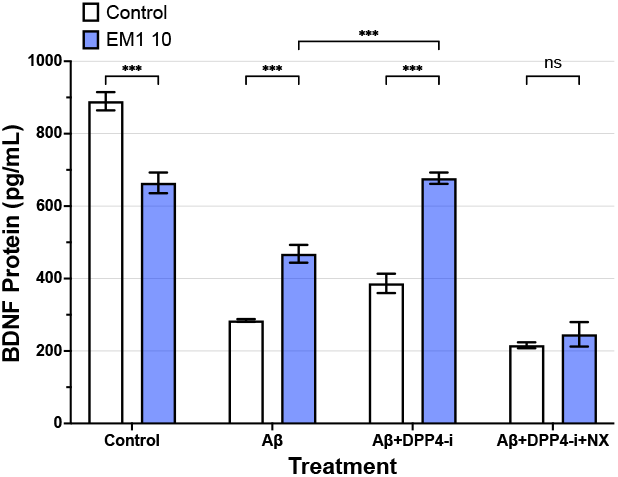
Effect of Sitagliptin, EM1, and EM2 on BDNF Concentration (*F*_3,16_ = 141.8)

## 3 Discussion

### 3.1 MOR Activation: Detrimental in Healthy Cells, Beneficial in A*β*_42_ Treated Cells

Findings in the first phase of this study showed that MOR activation through the endogenous EM1 and EM2 mu-opioid agonists increased neuronal SK-N-SH cell survival when exposed to A*β*_42_ in a dose-dependent manner. However, MOR activation decreased neuronal SK-N-SH cell survival in healthy cells in the inverse-dose-dependent manner. In this case, adding EM1 and EM2 in healthy cells disrupts the delicate balance of MOR activation and inhibition, but exposure to A*β*_42_ downregulates this MOR activation balance. This may explain why MOR activation through EM1 and EM2 is protective in A*β*_42_ exposed cells – however, the mechanism through which MOR activation improves cell vitality requires additional research.

The inhibition of MOR through NX, a known MOR antagonist, was similarly detrimental in healthy neuronal SK-N-SH cells. It is likely that MOR inhibition by NX also disrupt the delicate MOR activation balance. However, MOR inhibition by NX pretreatment followed by EM1 treatment prevented any increase in cell survival and led to survival rates lower compared to cells treated with only A*β*_42_. This supports the hypothesis that MOR activation is necessary for EM1’s protective effects at 1 μM and 10 μM concentrations.

Enormous efforts have been underway to develop an effective treatment for AD. Recent advances in AD research have suggested the potential of opioid receptors as a major regulator of memory processes, especially the mu-opioid receptor. However, numerous studies show conflicting results: it is unclear whether MOR activation improves or impairs memory processes. This study demonstrated the protective nature of MOR activation in A*β*_42_ treated cells and the detrimental effect of MOR activation in healthy cells. It is plausible that this discrepancy results from the specific type of A*β* peptide: A*β*_42_ induces MOR downregulation, while the relatively less toxic A*β*_40_ does not induce MOR downregulation.

### 3.2 High Concentrations of EM1 and EM2 Treatment Protect Against Oxidative Damage but Not Against Neuroinflammation

Treatment of EM1 and EM2 at higher concentrations of 10 μM and 20 μM resulted in the greatest improvement in cell survival rates in A*β*_42_ exposed cells. Based on previous research, concentrations of 5 μM of EM1 and EM2 are likely to induce the maximum capacity of MOR activation [75]. This may explain why the addition of 10 μM and 20 μM of EM1 and EM2 substantially improved in cell survival rates. I hypothesize that after reaching maximum MOR activation capacity, excess intracellular EM1 and EM2 may be able to directly interact with the intracellular A*β*_42_. One possible mechanism through which EM1 and EM2 may directly interact with intracellular A*β*_42_ is by mitigating the deleterious effects of A*β*_42_ generated reactive oxygen species, which results in oxidative damage. EM1 and EM2 at high concentrations (10 μM) significantly improved survival rates of cells exposed to rotenone, a well documented molecule for inducing oxidative stress through reactive oxygen species generation similar to A*β*_42_. This supports previous evidence that EM1 and EM2 may play an antioxidative and free radical scavenging role in the brain and thus provide protection against free radicals commonly found in many neurodegenerative disorders [45, 76]. This study also demonstrates that EM1 and EM2 treatment had no significant effect on cell survival in a LPS model of neuroinflammation, whether through MOR activation or potential intracellular interactions.

## 4 Discussion

Phase one of this study discovered that the MOR is likely to be downregulated in AD, and subsequent MOR activation through EM1 and EM2 improved SK-N-SH neuronal cell survival in AD. However, findings from phase one revealed the possibility of EM1 and EM2 mediating its effects outside of MOR activation. Subsequently, phase two of this study focused on determining the extent to which intracellular EM1 and EM2 could interact with A*β*_42_ plaques and whether preventing the degradation of intracellular EM1 and EM2 could enhance its effects.

### 4.1 Sitagliptin Is Able to Interact With EM1 and EM2

Although some studies have shown that DPP4 is co-localized with endomorphins in specific brain regions [59], relatively few studies have explored the interplay between DPP4, EM1, and EM2 in neuronal cells expressing aggregated A*β*_42_ plaques.

Sitagliptin, a drug that inhibits DPP4 [77], demonstrated a similar binding structure and affinity with EM1. Additionally, based on the high accuracy of docking and the number of hydrogen bonds and pi-pi stacked bonds [78] of EM1 to DPP4, it is highly likely that DPP4 can interact and therefore degrade EM1. Thus, DPP4 inhibition can prolong the half-life of intracellular EM1. The similar structures of EM1 and EM2 indicate that the half-life of intracellular EM2 can also be prolonged.

### 4.2 Sitagliptin Mediated DPP4-Inhibition Enhances the Effects of EM1 and EM2

The present study showed that high concentrations of EM1 and EM2 treatment (10 μM) with sitagliptin increased both the negative and positive effects of EM1 and EM2.

Interestingly, low concentrations of EM1 and EM2 treatment (1 μM) with sitagliptin had minimal or insignificant effects. Phase one of this study found that maximum MOR activation only occurs at or around 5 μM of EM1 or EM2 treatment for SK-N-SH neuronal cells. Thus, at low concentrations of 1 μM of EM1 or EM2 treatment, all endomorphins are likely to be bound to the MOR binding domain. As a result, DPP4 does not interact with the bounded EM1 and EM2. On the other hand, DPP4 is likely to exert a significant interaction with unbounded intracellular EM1 and EM2, such as when maximum MOR activation is reached after the excess addition of 10 μM of EM1 or EM2.

Since the treatment of 10 μM of EM1 and EM2 with sitagliptin increased cell survival compared to the treatment of 10 μM of EM1 and EM2 alone, increasing the half-lives of intracellular EM1 and EM2 by preventing DPP4-mediated degradation is protective in cells exposed to A*β*_42_. These results may explain why DPP4 inhibition can mitigate the detrimental effects associated with AD in previous rat and human studies [58, 60, 77, 79–82].

### 4.3 Intracellular EM1 Interaction, Not MOR Activation, Prevents A*β*_42_ Aggregation

To date, only one study has researched the effects of EM2 on restricting A*β*_42_ aggregation [53]. Our research demonstrates that intracellular EM1 significantly reduces A*β*_42_ aggregation, with combined sitagliptin and EM1 treatment further enhancing this effect. Notably, NX pretreatment had no discernible effect on A*β*_42_ concentration compared to EM1 treatment alone, suggesting that intracellular EM1—rather than MOR activation—is primarily responsible for limiting A*β*_42_ aggregation.

Based on these findings, we hypothesize that EM1 may function as an intracellular A*β*-breaker molecule. Structural analysis of EM1 reveals a tryptophan residue at the third position, and previous research has shown that tryptophan, an aromatic amino acid, has the greatest amyloidogenic propensity among amino acids [3]. This suggests that the tryptophan in EM1 may play a critical role in inhibiting A*β*_42_ aggregation through competitive binding mechanisms. However, additional studies are needed to confirm EM1’s role as a *β*-breaker molecule, including multiple ELISAs for smaller A*β*_1−40_ and A*β*_1−36_ peptides to determine whether EM1 can degrade longer A*β*_42_ into less toxic fragments [19, 83].

The *β*-breaker potential of EM1 in turn limits the toxicity exerted by aggregated A*β* plaques, including apoptosis activity and *H*_2_*O*_2_ free radical concentrations in cells. The decreased A*β*_42_ concentrations seem to play a role in decreasing caspase–3/7 activity, an indirect indicator of cell apoptosis activity, and *H*_2_*O*_2_ concentration, a metabolic by-product of reactive oxygen species that is a source of oxidative stress. 10 μM of EM1 treatment decreased caspase activity and *H*_2_*O*_2_ concentrations while sitagliptin and EM1 treatment further decreased caspase activity and *H*_2_*O*_2_ concentrations. However, NX pretreatment did not significantly affect caspase activity and *H*_2_*O*_2_ concentration. Therefore, it is likely that either the addition of 10 μM intracellular EM1 directly lowers cell apoptosis and *H*_2_*O*_2_ generation through an unknown pathway, or that the decreased aggregation of A*β*_42_ as a result of intracellular EM1 interaction subsequently reduced A*β*_42_ caused caspase activity and *H*_2_*O*_2_ generation.

### 4.4 MOR Activation, Not Intracellular EM1 Interaction, Increases Neurite Outgrowth and BDNF Concentrations

As EM1 has been found in abundance within the hippocampal CA1, CA2, and CA3 regions [7, 14, 33, 34], phase two of this study also examined the effects of EM1 on the hippocampal neurite outgrowth process. The addition of 10 μM of EM1 with sitagliptin did increase the total number of neurite intersections compared to 10 μM of EM1 treatment alone, but NX treatment reduced the total number of neurite intersections. These results are mirrored in BDNF protein concentrations. Therefore, it is likely that intracellular EM1 has no effect on the process of neurite growth. Rather, this is the first study to associate MOR activation in an AD cell model with increased BDNF concentrations, which in turn induces neurite outgrowth. Though these results support a previous study on increased BDNF concentrations following the activation of the *δ*-opioid receptor [84], no other study has researched *μ*-opioid receptor activation and BDNF concentrations in an AD model. Thus, further studies are needed to confirm my results. As DPP4 inhibition also increases GLP-1 and GLP-2 levels, future research could determine the extent to which GLP-1 and GLP-2 promotes BDNF protein concentrations.

Finally, this study might explain the decreased BDNF concentrations found in AD patients [85]. The first phase of this study demonstrated the likelihood of MOR downregulation in AD. As MOR activation increased BDNF protein concentrations in an AD cell model, it is plausible that MOR downregulation leads to the reduced BDNF concentrations in AD patients.

## 5 Conclusion

This study reveals several important implications for understanding the role of opioid systems in Alzheimer’s disease: 1) MOR signaling is tightly regulated in healthy neurons; 2) MOR becomes downregulated in AD, and its activation through EM1 and EM2 confers dose-dependent protection against A*β*_42_ toxicity, particularly by mitigating oxidative stress; 3) MOR activation promotes neurite outgrowth and increases BDNF expression, suggesting direct involvement in preserving memory circuits; 4) Intracellular EM1, when protected from degradation by DPP4 inhibition, effectively reduces A*β*_42_ aggregation and decreases *H*_2_*O*_2_ concentrations; and 5) MOR activation upregulates the ACE2 receptor pathway, potentially counteracting its downregulation in AD.

These findings have significant therapeutic implications, suggesting that dual-targeting approaches focusing on both MOR activation and intracellular EM1 stabilization could provide comprehensive neuroprotection in AD. The use of DPP4 inhibitors like sitagliptin, which are already approved for treating diabetes, represents a potential avenue for repurposing existing medications for AD treatment.

Future studies should investigate whether the amplified protective effects of DPP4 inhibitors combined with EM1 or EM2 involve insulin or GLP-1 signaling pathways. Additionally, further characterization of endomorphins’ antioxidant and *β*-breaker potential could lead to the development of novel therapeutic agents that specifically target A*β* aggregation while preserving critical memory processes.

## Acknowledgments

I would like to express my gratitude towards Professor Zhu, who has supported and mentored me for the entire 4 years of this research project. I would also like to thank Dr. Weseley for her generous advice and recommendations over my project.

## Notes

### Competing Interest Statement

The authors have declared no competing interest.

## References

[1] Colin L. Masters et al. “Alzheimer’s Disease”. In: Nature Reviews Disease Primers 1.1 (1 Oct. 15, 2015), pp. 1–18. issn: 2056-676X. doi: 10.1038/nrdp.2015.56. url: https://www.nature.com/articles/nrdp201556 (visited on 08/19/2022).

[2] Rudy J. Castellani, Raj K. Rolston, and Mark A. Smith. “Alzheimer Disease”. In: Disease-a-Month. Alzheimer Disease 56.9 (Sept. 1, 2010), pp. 484–546. issn: 0011-5029. doi: 10.1016/j.disamonth.2010.06.001. url: https://www.sciencedirect.com/science/article/pii/S0011502910000684 (visited on 08/19/2022).

[3] Anat Frydman-Marom et al. “Structural Basis for Inhibiting β-Amyloid Oligomerization by a Non-coded β-Breaker-Substituted Endomorphin Analogue”. In: ACS Chemical Biology 6.11 (Nov. 18, 2011), pp. 1265–1276. issn: 1554-8929. doi: 10.1021/cb200103h. url: https://doi.org/10.1021/cb200103h (visited on 08/18/2022).

[4] Said AbdAlla et al. “ACE Inhibition with Captopril Retards the Development of Signs of Neurodegeneration in an Animal Model of Alzheimer’s Disease”. In: International Journal of Molecular Sciences 14.8 (8 Aug. 2013), pp. 16917–16942. issn: 1422-0067. doi: 10.3390/ijms140816917. url: https://www.mdpi.com/1422-0067/14/8/16917 (visited on 08/18/2022).

[5] Efthalia Angelopoulou and Christina Piperi. “DPP-4 Inhibitors: A Promising Therapeutic Approach against Alzheimer’s Disease”. In: Annals of Translational Medicine 6.12 (June 2018), p. 255. issn: 2305-5839. doi: 10.21037/atm.2018.04.41. pmid: 30069457.

[6] Verena H. Finder. “Alzheimer’s Disease: A General Introduction and Pathomechanism”. In: Journal of Alzheimer’s disease: JAD 22 Suppl 3 (2010), pp. 5–19. issn: 1875-8908. doi: 10.3233/JAD-2010-100975. pmid: 20858960.

[7] Zhiyou Cai and Anna Ratka. “Opioid System and Alzheimer’s Disease”. In: Neuromolecular Medicine 14.2 (June 2012), pp. 91–111. issn: 1559-1174. doi: 10.1007/s12017-012-8180-3. pmid: 22527793.

[8] Vikas Dhikav and Kuljeet Anand. “Potential Predictors of Hippocampal Atrophy in Alzheimer’s Disease”. In: Drugs & Aging 28.1 (Jan. 1, 2011), pp. 1–11. issn: 1179-1969. doi: 10.2165/11586390-000000000-00000. pmid: 21174483.

[9] Kwasi G. Mawuenyega et al. “Decreased Clearance of Cns Beta Amyloid in Alzheimer s Disease”. In: Science (2010). doi: 10.1126/science.1197623. pmid: 21148344.

[10] Masafumi Sakono and Tamotsu Zako. “Amyloid Oligomers: Formation and Toxicity of Abeta Oligomers”. In: The FEBS journal 277.6 (Mar. 2010), pp. 1348–1358. issn: 1742-4658. doi: 10.1111/j.1742-4658.2010.07568.x. pmid: 20148964.

[11] Jason Weller and Andrew Budson. “Current Understanding of Alzheimer’s Disease Diagnosis and Treatment”. In: F1000Research 7 (July 31, 2018), F1000 Faculty Rev–1161. issn: 2046-1402. doi: 10.12688/f1000research.14506.1. pmid: 30135715. url: https://www.ncbi.nlm.nih.gov/pmc/articles/PMC6073093/ (visited on 08/24/2022).

[12] Jitin Bali et al. “Cellular Basis of Alzheimer’s Disease”. In: Annals of Indian Academy of Neurology 13 (Suppl2 Dec. 2010), S89–S93. issn: 0972-2327. doi: 10.4103/0972-2327.74251. pmid: 21369424. url: https://www.ncbi.nlm.nih.gov/pmc/articles/PMC3039159/ (visited on 08/24/2022).

[13] Mohammad Reza Zarrindast, Majid Navaeian, and Mohammad Nasehi. “Influence of Three-Day Morphine-Treatment upon Impairment of Memory Consolidation Induced by Cannabinoid Infused into the Dorsal Hippocampus in Rats”. In: Neuroscience Research 69.1 (Jan. 1, 2011), pp. 51–59. issn: 0168-0102. doi: 10.1016/j.neures.2010.09.007. url: https://www.sciencedirect.com/science/article/pii/S0168010210028063 (visited on 07/05/2022).

[14] L. Jamot et al. “Differential Involvement of the Mu and Kappa Opioid Receptors in Spatial Learning”. In: Genes, Brain and Behavior 2.2 (2003), pp. 80–92. issn: 1601-183X. doi: 10.1034/j.1601-183X.2003.00013.x. url: https://onlinelibrary.wiley.com/doi/abs/10.1034/j.1601-183X.2003.00013.x(visited on 08/19/2022).

[15] Michele L. Simmons and Charles Chavkin. “Endogenous Opioid Regulation of Hippocampal Function”. In: International Review of Neurobiology. Ed. by Ronald J. Bradley, R. Adron Harris, and Peter Jenner. Vol. 39. Academic Press, Jan. 1, 1996, pp. 145–196. doi: 10.1016/S0074-7742(08)60666-2. url: https://www.sciencedirect.com/science/article/pii/S0074774208606662 (visited on 08/19/2022).

[16] Sheryl Martin-Schild et al. “Differential Distribution of Endomorphin 1- and Endomorphin 2-like Immunoreactivities in the CNS of the Rodent.” In: The Journal of Comparative Neurology 405.4 (Mar. 22, 1999), pp. 450–471. doi: 10.1002/(SICI)1096-9861(19990322)405:4<450::AID-CNE2>3.0.CO;2-%23. pmid: 10098939.

[17] Esraa Mohamed et al. “Endogenous Opioid Peptides and Brain Development: Endomorphin-1 and Nociceptin Play a Sex-Specific Role in the Control of Oligodendrocyte Maturation and Brain Myelination”. In: Glia 68.7 (July 2020), pp. 1513–1530. issn: 1098-1136. doi: 10.1002/glia.23799. pmid: 32065429.

[18] Fred Nyberg. “Opioid Peptides in Cerebrospinal Fluid-Methods for Analysis and Their Significance in the Clinical Perspective”. In: Frontiers in Bioscience: A Journal and Virtual Library 9 (Sept. 1, 2004), pp. 3510–3525. issn: 1093-9946. doi: 10.2741/1497. pmid: 15353373.

[19] Maria Laura Giuffrida et al. “Beta-Amyloid Monomers Are Neuroprotective”. In: The Journal of Neuroscience: The Official Journal of the Society for Neuroscience 29.34 (Aug. 26, 2009), pp. 10582–10587. issn: 1529-2401. doi: 10.1523/JNEUROSCI.1736-09.2009. pmid: 19710311.

[20] Shu-Hui Xin et al. “Clearance of Amyloid Beta and Tau in Alzheimer’s Disease: From Mechanisms to Therapy”. In: Neurotoxicity Research 34.3 (Oct. 2018), pp. 733–748. issn: 1476-3524. doi: 10.1007/s12640-018-9895-1. pmid: 29626319.

[21] David J. Bonda et al. “Oxidative Stress in Alzheimer Disease: A Possibility for Prevention”. In: Neuropharmacology 59.4-5 (2010), pp. 290–294. issn: 1873-7064. doi: 10.1016/j.neuropharm.2010.04.005. pmid: 20394761.

[22] C. Cheignon et al. “Oxidative Stress and the Amyloid Beta Peptide in Alzheimer’s Disease”. In: Redox Biology 14 (Apr. 1, 2018), pp. 450–464. issn: 2213-2317. doi: 10.1016/j.redox.2017.10.014. url: https://www.sciencedirect.com/science/article/pii/S2213231717307267 (visited on 08/28/2022).

[23] Zhichun Chen and Chunjiu Zhong. “Oxidative Stress in Alzheimer’s Disease”. In: Neuroscience Bulletin 30.2 (Apr. 2014), pp. 271–281. issn: 1995-8218. doi: 10.1007/s12264-013-1423-y. pmid: 24664866.

[24] Jeffrey A. Klein and Susan L. Ackerman. “Oxidative Stress, Cell Cycle, and Neurodegeneration”. In: Journal of Clinical Investigation 111.6 (Mar. 15, 2003), pp. 785–793. issn: 0021-9738. doi: 10.1172/JCI200318182. pmid: 12639981. url: https://www.ncbi.nlm.nih.gov/pmc/articles/PMC153779/ (visited on 08/18/2022).

[25] Maria Lehtinen and Azad Bonni. “Modeling Oxidative Stress in the Central Nervous System”. In: Current Molecular Medicine 6.8 (Dec. 1, 2006), pp. 871–881. issn: 15665240. doi: 10.2174/156652406779010786. url: http://www.eurekaselect.com/openurl/content.php?genre=article&issn=1566-5240&volume=6&issue=8&spage=871 (visited on 08/18/2022).

[26] Xinkun Wang and Elias Michaelis. “Selective Neuronal Vulnerability to Oxidative Stress in the Brain”. In: Frontiers in Aging Neuroscience 2 (2010). ISSN: 1663-4365. doi: 10.3389/fnagi.2010.00012. url: https://www.frontiersin.org/articles/10.3389/fnagi.2010.00012 (visited on 08/18/2022).

[27] Christian Behl and Bernd Moosmann. “Antioxidant Neuroprotection in Alzheimer’s Disease as Preventive and Therapeutic Approach2 2This Article Is Part of a Series of Reviews on “Causes and Consequences of Oxidative Stress in Alzheimer’s Disease.” The Full List of Papers May Be Found on the Homepage of the Journal.” In: Free Radical Biology and Medicine 33.2 (July 15, 2002), pp. 182–191. issn: 0891-5849. doi: 10.1016/S0891-5849(02)00883-3. url: https://www.sciencedirect.com/science/article/pii/S0891584902008833 (visited on 08/28/2022).

[28] Bernd Moosmann and Christian Behl. “Antioxidants as Treatment for Neurodegenerative Disorders”. In: Expert Opinion on Investigational Drugs 11.10 (Oct. 2002), pp. 1407–1435. issn: 1354-3784. doi: 10.1517/13543784.11.10.1407. pmid: 12387703.

[29] Chunyan Guo et al. “Oxidative Stress, Mitochondrial Damage and Neurodegenerative Diseases”. In: Neural Regeneration Research 8.21 (July 25, 2013), pp. 2003–2014. issn: 1673-5374. doi: 10.3969/j.issn.1673-5374.2013.21.009. pmid: 25206509. url: https://www.ncbi.nlm.nih.gov/pmc/articles/PMC4145906/ (visited on 08/24/2022).

[30] David S. Jessop et al. “Novel Opioid Peptides Endomorphin-1 and Endomorphin-2 Are Present in Mammalian Immune Tissues”. In: Journal of Neuroimmunology 106.1 (July 1, 2000), pp. 53–59. issn: 0165-5728, 1872-8421. doi: 10.1016/S0165-5728(99)00216-7. pmid: 10814782. url: https://www.jni-journal.com/article/S0165-5728(99)00216-7/fulltext (visited on 07/14/2022).

[31] Maria Waldhoer, Selena E. Bartlett, and Jennifer L. Whistler. “Opioid Receptors”. In: Annual Review of Biochemistry 73.1 (2004), pp. 953–990. doi: 10.1146/annurev.biochem.73.011303.073940. pmid: 15189164. url: https://doi.org/10.1146/annurev.biochem.73.011303.073940 (visited on 08/24/2022).

[32] Gavril W. Pasternak and Ying-Xian Pan. “Mu Opioids and Their Receptors: Evolution of a Concept”. In: Pharmacological Reviews 65.4 (Oct. 1, 2013). Ed. by David R. Sibley, pp. 1257–1317. issn: 0031-6997, 1521-0081. doi: 10.1124/pr.112.007138. pmid: 24076545. url: https://pharmrev.aspetjournals.org/content/65/4/1257 (visited on 07/09/2022).

[33] Eric Erbs et al. “A Mu–Delta Opioid Receptor Brain Atlas Reveals Neuronal Co-Occurrence in Subcortical Networks”. In: Brain Structure & Function 220.2 (Mar. 1, 2015), pp. 677–702. doi: 10.1007/s00429-014-0717-9. pmid: 24623156.

[34] Carrie T. Drake and Teresa A. Milner. “Mu Opioid Receptors Are in Somatodendritic and Axonal Compartments of GABAergic Neurons in Rat Hippocampal Formation”. In: Brain Research 849.1 (Dec. 4, 1999), pp. 203–215. issn: 0006-8993. doi: 10.1016/S0006-8993(99)01910-1. url: https://www.sciencedirect.com/science/article/pii/S0006899399019101 (visited on 08/20/2022).

[35] Min-Ho Nam et al. “Expression of M -Opioid Receptor in CA1 Hippocampal Astrocytes”. In: Experimental Neurobiology 27.2 (Apr. 2018), pp. 120–128. issn: 1226-2560. doi: 10.5607/en.2018.27.2.120. pmid: 29731678.

[36] Anjana Bali, Puneet Kaur Randhawa, and Amteshwar Singh Jaggi. “Interplay between RAS and Opioids: Opening the Pandora of Complexities”. In: Neuropeptides 48.4 (Aug. 2014), pp. 249–256. issn: 1532-2785. doi: 10.1016/j.npep.2014.05.002. pmid: 24877897.

[37] Gerald D. Hess, James A. Joseph, and George S. Roth. “Effect of Age on Sensitivity to Pain and Brain Opiate Receptors”. In: Neurobiology of Aging 2.1 (Mar. 1, 1981), pp. 49–55. issn: 0197-4580. doi: 10.1016/0197-4580(81)90059-2. url: https://www.sciencedirect.com/science/article/pii/0197458081900592 (visited on 08/20/2022).

[38] J. M. Hiller, Y. Itzhak, and E. J. Simon. “Selective Changes in Mu, Delta and Kappa Opioid Receptor Binding in Certain Limbic Regions of the Brain in Alzheimer’s Disease Patients”. In: Brain Research 406.1-2 (Mar. 17, 1987), pp. 17–23. issn: 0006-8993. doi: 10.1016/0006-8993(87)90764-5. pmid: 3032356.

[39] Anne-Marie Mathieu-Kia et al. “μ-, δ- and κ-Opioid Receptor Populations Are Differentially Altered in Distinct Areas of Postmortem Brains of Alzheimer’s Disease Patients”. In: Brain Research 893.1 (Mar. 2, 2001), pp. 121–134. issn: 0006-8993. doi: 10.1016/S0006-8993(00)03302-3. url: https://www.sciencedirect.com/science/article/pii/S0006899300033023 (visited on 08/26/2022).

[40] Jon-Kar Zubieta, Robert F. Dannals, and J. James Frost. “Gender and Age Influences on Human Brain Mu-Opioid Receptor Binding Measured by PET”. In: American Journal of Psychiatry 156.6 (June 1999), pp. 842–848. issn: 0002-953X. doi: 10.1176/ajp.156.6.842. url: https://ajp.psychiatryonline.org/doi/full/10.1176/ajp.156.6.842 (visited on 08/20/2022).

[41] Yuan Feng et al. “Current Research on Opioid Receptor Function”. In: Current Drug Targets 13.2 (Feb. 2012), pp. 230–246. issn: 1873-5592. doi: 10.2174/138945012799201612. pmid: 22204322.

[42] Ream Al Hasani and Michael R. Bruchas. “Molecular Mechanisms of Opioid Receptor Dependent Signaling and Behavior”. In: Anesthesiology 115.6 (2011), pp. 1363–1381. issn: 0003-3022. doi: 10.1097/ALN.0b013e318238bba6. pmid: 22020140. url: https://www.ncbi.nlm.nih.gov/pmc/articles/PMC3698859/ (visited on 07/12/2022).

[43] Jakub Fichna et al. “The Endomorphin System and Its Evolving Neurophysiological Role”. In: Pharmacological Reviews 59.1 (Mar. 1, 2007), pp. 88–123. doi: 10.1124/pr.59.1.3. pmid: 17329549.

[44] G. Horvath. “Endomorphin-1 and Endomorphin-2: Pharmacology of the Selective Endogenous Mu-Opioid Receptor Agonists”. In: Pharmacology & Therapeutics 88.3 (Dec. 2000), pp. 437–463. issn: 0163-7258. doi: 10.1016/s0163-7258(00)00100-5. pmid: 11337033.

[45] Xin Lin et al. “Protective Effects of Endomorphins, Endogenous Opioid Peptides in the Brain, on Human Low Density Lipoprotein Oxidation”. In: The FEBS journal 273.6 (Mar. 2006), pp. 1275–1284. issn: 1742-464X. doi: 10.1111/j.1742-4658.2006.05150.x. pmid: 16519691.

[46] Claudio Castellano et al. “The Effects of Morphine on Memory Consolidation in Mice Involve Both D1 and D2 Dopamine Receptors”. In: Behavioral and Neural Biology 61.2 (Mar. 1, 1994), pp. 156–161. issn: 0163-1047. doi: 10.1016/S0163-1047(05)80069-X. url: https://www.sciencedirect.com/science/article/pii/S016310470580069X (visited on 07/05/2022).

[47] Yun Feng et al. “Endomorphins and Morphine Limit Anoxia-Reoxygenation-Induced Brain Mitochondrial Dysfunction in the Mouse”. In: Life Sciences 82.13-14 (Mar. 26, 2008), pp. 752–763. issn: 0024-3205. doi: 10.1016/j.lfs.2008.01.004. pmid: 18272183.

[48] Lynn Nadel and Oliver Hardt. “Update on Memory Systems and Processes”. In: Neuropsychopharmacology 36.1 (Jan. 2011), pp. 251–273. issn: 0893-133X. doi: 10.1038/npp.2010.169. pmid: 20861829. url: https://www.ncbi.nlm.nih.gov/pmc/articles/PMC3055510/ (visited on 07/21/2022).

[49] Ramasamy Perumal and I. Tan. “What Proteins Are Involved in Learning and Memory?” In: IUBMB Life 59.7 (2007), pp. 465–468. ISSN: 1521-6551. doi: 10.1080/15216540601055331. url: https://onlinelibrary.wiley.com/doi/abs/10.1080/15216540601055331 (visited on 07/16/2022).

[50] Y. Shiigi, M. Takahashi, and H. Kaneto. “Facilitation of Memory Retrieval by Pretest Morphine Mediated by Mu but Not Delta and Kappa Opioid Receptors”. In: Psychopharmacology 102.3 (1990), pp. 329–332. issn: 0033-3158. doi: 10.1007/BF02244099. pmid: 2174567.

[51] Choon-Gon Jang et al. “Impaired Water Maze Learning Performance in μ-Opioid Receptor Knockout Mice”. In: Molecular Brain Research 117.1 (Sept. 10, 2003), pp. 68–72. issn: 0169-328X. doi: 10.1016/S0169-328X(03)00291-2. url: https://www.sciencedirect.com/science/article/pii/S0169328X03002912 (visited on 07/14/2022).

[52] J. E. Zadina et al. “A Potent and Selective Endogenous Agonist for the Mu-Opiate Receptor”. In: Nature 386.6624 (Apr. 3, 1997), pp. 499–502. issn: 0028-0836. doi: 10.1038/386499a0. pmid: 9087409.

[53] Viktor Szegedi. “Endomorphin-2, an Endogenous Tetrapeptide, Protects against Aβ1-42 in Vitro and in Vivo”. In: (2006). doi: 10.1096/fj.05-4891fje. url: https://faseb.onlinelibrary.wiley.com/doi/10.1096/fj.05-4891fje (visited on 08/18/2022).

[54] Ghaith Al-Badri et al. “Tackling Dipeptidyl Peptidase IV in Neurological Disorders”. In: Neural Regeneration Research 13.1 (Jan. 2018), pp. 26–34. issn: 1673-5374. doi: 10.4103/1673-5374.224365. pmid: 29451201.

[55] Nian Gong et al. “Activation of Spinal Glucagon-like Peptide-1 Receptors Specifically Suppresses Pain Hypersensitivity”. In: The Journal of Neuroscience: The Official Journal of the Society for Neuroscience 34.15 (Apr. 9, 2014), pp. 5322–5334. issn: 1529-2401. doi: 10.1523/JNEUROSCI.4703-13.2014. pmid: 24719110.

[56] Eriko Komiya et al. “Peripheral Endomorphins Drive Mechanical Alloknesis under the Enzymatic Control of CD26/DPPIV”. In: The Journal of Allergy and Clinical Immunology 149.3 (Mar. 2022), pp. 1085–1096. issn: 1097-6825. doi: 10.1016/j.jaci.2021.08.003. pmid: 34411589.

[57] Randi Shane, Sherwin Wilk, and Richard J. Bodnar. “Modulation of Endomorphin-2-Induced Analgesia by Dipeptidyl Peptidase IV.” In: Brain Research 815.2 (Jan. 9, 1999), pp. 278–286. doi: 10.1016/s0006-8993(98)01121-4. pmid: 9878785.

[58] Nehru Sai Suresh Chalichem, Pindiprolu S. S. Sai Kiran, and Duraiswamy Basavan. “Possible Role of DPP4 Inhibitors to Promote Hippocampal Neurogenesis in Alzheimer’s Disease”. In: Journal of Drug Targeting 26.8 (Sept. 2018), pp. 670–675. issn: 1029-2330. doi: 10.1080/1061186X.2018.1433682. pmid: 29378454.

[59] Marina Baršun et al. “Human Dipeptidyl Peptidase III Acts as a Post-Proline-Cleaving Enzyme on Endomorphins.” In: Biological Chemistry 388.3 (Jan. 1, 2007), pp. 343–348. doi: 10.1515/bc.2007.039. pmid: 17338643.

[60] Shuyi Chen et al. “DPP-4 Inhibitor Improves Learning and Memory Deficits and AD-like Neurodegeneration by Modulating the GLP-1 Signaling”. In: Neuropharmacology 157 (Oct. 2019), p. 107668. issn: 1873-7064. doi: 10.1016/j.neuropharm.2019.107668. pmid: 31199957.

[61] Luca Colucci-D’Amato, Luisa Speranza, and Floriana Volpicelli. “Neurotrophic Factor BDNF, Physiological Functions and Therapeutic Potential in Depression, Neurodegeneration and Brain Cancer”. In: International Journal of Molecular Sciences 21.20 (Oct. 21, 2020), p. 7777. ISSN: 1422-0067. doi:10.3390/ijms21207777. pmid: 33096634. url: https://www.ncbi.nlm.nih.gov/pmc/articles/PMC7589016/ (visited on 07/12/2022).

[62] Ewelina Palasz et al. “BDNF as a Promising Therapeutic Agent in Parkinson’s Disease”. In: International Journal of Molecular Sciences 21.3 (Feb. 10, 2020), p. 1170. ISSN: 1422-0067. doi: 10.3390/ijms21031170. pmid: 32050617. url: https://www.ncbi.nlm.nih.gov/pmc/articles/PMC7037114/ (visited on 07/12/2022).

[63] Devin K. Binder and Helen E. Scharfman. “Brain-Derived Neurotrophic Factor”. In: Growth factors (Chur, Switzerland) 22.3 (Sept. 2004), pp. 123–131. issn: 0897-7194. doi: 10.1080/08977190410001723308. pmid: 15518235. url: https://www.ncbi.nlm.nih.gov/pmc/articles/PMC2504526/ (visited on 07/12/2022).

[64] E. J. Huang and L. F. Reichardt. “Neurotrophins: Roles in Neuronal Development and Function”. In: Annual Review of Neuroscience 24 (2001), pp. 677–736. issn: 0147-006X. doi: 10.1146/annurev.neuro.24.1.677. pmid: 11520916.

[65] A. Acheson et al. “A BDNF Autocrine Loop in Adult Sensory Neurons Prevents Cell Death”. In: Nature 374.6521 (Mar. 30, 1995), pp. 450–453. issn: 0028-0836. doi: 10.1038/374450a0. pmid: 7700353.

[66] Francesca Stanzione, Ilenia Giangreco, and Jason C. Cole. “Use of Molecular Docking Computational Tools in Drug Discovery”. In: Progress in Medicinal Chemistry 60 (2021), pp. 273–343. issn: 0079-6468. doi: 10.1016/bs.pmch.2021.01.004. pmid: 34147204.

[67] Luca Pinzi and Giulio Rastelli. “Molecular Docking: Shifting Paradigms in Drug Discovery”. In: International Journal of Molecular Sciences 20.18 (Sept. 4, 2019), E4331. issn: 1422-0067. doi: 10.3390/ijms20184331. pmid: 31487867.

[68] Anna E. Lohning et al. “A Practical Guide to Molecular Docking and Homology Modelling for Medicinal Chemists”. In: Current Topics in Medicinal Chemistry 17.18 (2017), pp. 2023–2040. issn: 1873-4294. doi: 10.2174/1568026617666170130110827. pmid: 28137238.

[69] Oleg Trott and Arthur J. Olson. “AutoDock Vina: Improving the Speed and Accuracy of Docking with a New Scoring Function, Efficient Optimization, and Multithreading”. In: Journal of Computational Chemistry 31.2 (Jan. 30, 2010), pp. 455–461. issn: 1096-987X. doi: 10.1002/jcc.21334. pmid: 19499576.

[70] Sargis Dallakyan and Arthur J. Olson. “Small-Molecule Library Screening by Docking with PyRx”. In: Methods in Molecular Biology (Clifton, N.J.) 1263 (2015), pp. 243–250. issn: 1940-6029. doi: 10.1007/978-1-4939-2269-7_19. pmid: 25618350.

[71] Johan van Meerloo, Gertjan J. L. Kaspers, and Jacqueline Cloos. “Cell Sensitivity Assays: The MTT Assay”. In: Methods in Molecular Biology (Clifton, N.J.) 731 (2011), pp. 237–245. issn: 1940-6029. doi: 10.1007/978-1-61779-080-5_20. pmid: 21516412.

[72] Rosa A. Carrasco, Nancy B. Stamm, and Bharvin K. R. Patel. “One-Step Cellular Caspase-3/7 Assay”. In: BioTechniques 34.5 (May 2003), pp. 1064–1067. issn: 0736-6205. doi: 10.2144/03345dd02. pmid: 12765032.

[73] D. Oda et al. “H2O2 Oxidative Damage in Cultured Oral Epithelial Cells: The Effect of Short-Term Vitamin C Exposure”. In: Anticancer Research 21 (4A 2001), pp. 2719–2724. issn: 0250-7005. pmid: 11724346.

[74] Danielle E. Levitt, Katherine A. Adler, and Liz Simon. “HEMA 3 Staining: A Simple Alternative for the Assessment of Myoblast Differentiation”. In: Current Protocols in Stem Cell Biology 51.1 (Dec. 2019), e101. issn: 1938-8969. doi: 10.1002/cpsc.101. pmid: 31756292.

[75] G. Ponterio et al. “Powerful Inhibitory Action of Mu Opioid Receptors (MOR) on Cholinergic Interneuron Excitability in the Dorsal Striatum”. In: Neuropharmacology 75 (Dec. 1, 2013), pp. 78–85. issn: 0028-3908. doi: 10.1016/j.neuropharm.2013.07.006. url: https://www.sciencedirect.com/science/article/pii/S0028390813003237 (visited on 08/20/2022).

[76] Xin Lin et al. “Endomorphins, Endogenous Opioid Peptides, Provide Antioxidant Defense in the Brain against Free Radical-Induced Damage”. In: Biochimica et Biophysica Acta (BBA) - Molecular Basis of Disease 1639.3 (Nov. 20, 2003), pp. 195–202. issn: 0925-4439. doi: 10.1016/j.bbadis.2003.09.007. url: https://www.sciencedirect.com/science/article/pii/S0925443903001601 (visited on 08/19/2022).

[77] Ahmet Turan Isik et al. “The Effects of Sitagliptin, a DPP-4 Inhibitor, on Cognitive Functions in Elderly Diabetic Patients with or without Alzheimer’s Disease”. In: Diabetes Research and Clinical Practice 123 (Jan. 2017), pp. 192–198. issn: 1872-8227. doi: 10.1016/j.diabres.2016.12.010. pmid: 28056430.

[78] Steve Scheiner, Tapas Kar, and Jayasree Pattanayak. “Comparison of Various Types of Hydrogen Bonds Involving Aromatic Amino Acids”. In: Journal of the American Chemical Society 124.44 (Nov. 6, 2002), pp. 13257–13264. issn: 0002-7863. doi: 10.1021/ja027200q. pmid: 12405854.

[79] Jayasankar Kosaraju et al. “Linagliptin, a Dipeptidyl Peptidase-4 Inhibitor, Mitigates Cognitive Deficits and Pathology in the 3xTg-AD Mouse Model of Alzheimer’s Disease”. In: Molecular Neurobiology 54.8 (Oct. 2017), pp. 6074–6084. issn: 1559-1182. doi: 10.1007/s12035-016-0125-7. pmid: 27699599.

[80] Michele D’Amico et al. “Long-Term Inhibition of Dipeptidyl Peptidase-4 in Alzheimer’s Prone Mice”. In: Experimental Gerontology 45.3 (Mar. 2010), pp. 202–207. issn: 1873-6815. doi: 10.1016/j.exger.2009.12.004. pmid: 20005285.

[81] Michael Gejl et al. “In Alzheimer’s Disease, 6-Month Treatment with GLP-1 Analog Prevents Decline of Brain Glucose Metabolism: Randomized, Placebo-Controlled, Double-Blind Clinical Trial”. In: Frontiers in Aging Neuroscience 8 (2016), p. 108. issn: 1663-4365. doi: 10.3389/fnagi.2016.00108. pmid: 27252647.

[82] Christian Hölscher. “Central Effects of GLP-1: New Opportunities for Treatments of Neurodegenerative Diseases”. In: The Journal of Endocrinology 221.1 (Apr. 2014), T31–41. issn: 1479-6805. doi: 10.1530/JOE-13-0221. pmid: 23999914.

[83] Rakez Kayed and Cristian A. Lasagna-Reeves. “Molecular Mechanisms of Amyloid Oligomers Toxicity”. In: Journal of Alzheimer’s disease: JAD 33 Suppl 1 (2013), S67–78. issn: 1875-8908. doi: 10.3233/JAD-2012-129001. pmid: 22531422.

[84] Mary M. Torregrossa et al. “The Delta-Opioid Receptor Agonist (+)BW373U86 Regulates BDNF mRNA Expression in Rats”. In: Neuropsychopharmacology: Official Publication of the American College of Neuropsychopharmacology 29.4 (Apr. 2004), pp. 649–659. issn: 0893-133X. doi: 10.1038/sj.npp.1300345. pmid: 14647482.

[85] Jing-Hui Song, Jin-Tai Yu, and Lan Tan. “Brain-Derived Neurotrophic Factor in Alzheimer’s Disease: Risk, Mechanisms, and Therapy”. In: Molecular Neurobiology 52.3 (Dec. 2015), pp. 1477–1493. issn: 1559-1182. doi: 10.1007/s12035-014-8958-4. pmid: 25354497.

